# AAV-mediated MUC5AC siRNA delivery to prevent mucociliary dysfunction in asthma

**DOI:** 10.1101/2025.03.12.642720

**Authors:** Sahana Kumar, Maria Corkran, Yahya Cheema, Margaret A. Scull, Gregg A. Duncan

**Affiliations:** Department of Cell Biology & Molecular Genetics, Maryland Pathogen Research Institute (MPRI) University of Maryland, College Park, MD 20742; Fischell Department of Bioengineering, University of Maryland, College Park, MD 20742

## Abstract

The main structural components of mucus produced in the lung are mucin 5B (MUC5B) and mucin 5AC (MUC5AC) where a relatively higher expression of MUC5B is typical in health. In the lungs of individuals with asthma, there is a shift from MUC5B to MUC5AC as the predominantly secreted mucin which has been shown to impair mucociliary clearance (MCC) and increase mucus plug formation in the airways. Given its role in asthmatic lung disease, MUC5AC represents a potential therapeutic target where a gene delivery approach could be leveraged to modulate its expression. For these purposes, we explored adeno-associated virus serotype 6 (AAV6), as a lung-tropic viral gene vector to target airway epithelial cells and reduce MUC5AC expression via siRNA delivery. We confirmed that AAV6 was able to transduce epithelial cells in the airways of healthy mice with high transgene expression in mucus-secreting goblet cells. Using multiple particle tracking analysis, we observed that AAV6 was capable of penetrating both normal and MUC5AC-enriched mucus barriers. Successful transduction with AAV6 was also achieved in IL-13 stimulated human airway epithelial (HAE) cells differentiated at air-liquid interface (ALI). AAV6 expressing MUC5AC-targeting siRNA was evaluated as a prophylactic treatment in HAE cell cultures before IL-13 challenge. IL-13 stimulated HAE cultures treated with AAV6-MUC5AC siRNA had significantly reduced MUC5AC mRNA and protein expression compared to untreated controls. Mucociliary transport in IL-13 stimulated HAE cultures was also maintained and comparable to healthy controls following AAV6-MUC5AC siRNA treatment. Together, these findings support that AAV6 may be used as an inhaled gene therapy to suppress MUC5AC overexpression and restore normal airway clearance function in asthma.

## Introduction

Asthma is a chronic lung disease, affecting millions worldwide, adults and children alike^1^. Individuals with asthma typically experience coughing, difficulty breathing and wheezing which impacts their day to day activities and overall quality of life^2,3^. Allergic asthma is associated with airway inflammation, airway remodeling, and impaired clearance of mucus from the airway by coordinated beating of cilia on the epithelium, a process known as mucociliary clearance (MCC)^4,5^. As asthma worsens, excess mucus is produced leading to mucus plug formations due to impaired MCC^6^. Standard treatments for asthma, including corticosteroids and bronchoconstrictors, are focused primarily on managing the symptoms of inflammation and airway constriction. However, patients endure the disease for years and long-term use of these drugs can cause significant financial burden and have off-target effects^7–9^. Beyond over-the-counter expectorants which may provide temporary symptom relief, there has been minimal progress in therapeutic development to address muco-obstruction and airway plugging observed in severe asthma.

To achieve long-lasting therapeutic effects, gene therapy strategies have been developed and evaluated for their ability to inhibit or reduce specific Th2 inflammatory pathways in asthmatic airways. In addition, local delivery of gene therapies for asthma via inhaled or intranasal administration can enable less frequent dosing and reduce off-target effects^10^. So far, targeted gene therapeutic approaches for asthma have used both viral and non-viral vectors, loaded with either antisense oligonucleotides, siRNA, or mRNA that aim to antagonize or silence inflammatory mediators or shift the immune response from a Th2 response to a Th1 response^11–16^. In previous work, an adeno associated virus (AAV) encoding for receptor antagonist of IL-4RA was able to alleviate the Th2 allergic response in a mouse model of allergic asthma. This strategy also reduced mucus hypersecretion and airway hyperresponsiveness in ovalbumin sensitized mice^11^. Another study used AAV for sustained expression of IL-12 to enhance Th1 responses in OVA sensitized asthmatic mice where they observed reduced Th2 cytokines in treated mice^17^.

While inflammation and immune mediators are an important therapeutic target, fatal asthma attacks are often caused by mucus plugs which can partially or fully occlude small and large airways leading to restricted airflow^18–20^. Within the lung, mucus is formed by gel-forming mucin glycoproteins, mucin 5B (MUC5B) and mucin 5AC (MUC5AC) where each of these mucins possess distinct biochemical (e.g., glycosylation pattern) and biophysical properties (e.g., macromolecular assembly, crosslinking)^6^. Healthy airway mucus is comprised predominantly of MUC5B and with relatively lower levels of MUC5AC, while the contrary is observed in asthmatic airways, with MUC5AC being the predominant mucin^21^. Prior studies have shown that this change in mucus composition contributes to viscous mucus, impaired MCC, and mucus plug formations in airways^18,22,23^. These prior studies also established IL-13 mediated goblet cell hyperplasia in asthma leads to MUC5AC hypersecretion and pathological mucus production in asthma. A study by Siddiqui et al. targeted microRNA miR-141, which regulates IL-13-induced mucus secretion, and showed miR-141 disruption reduced goblet cell hyperplasia, MUC5AC expression, and total secreted mucus^24^. As such, MUC5AC warrants attention as a therapeutic target for treating asthma which could be addressed using a gene therapy strategy.

AAV is one of the leading viral vectors for gene therapy applications because it has broad tissue tropism and is non-pathogenic. Further, several AAV-based approaches have obtained FDA approval with many others in clinical trials^25,26^. To deliver MUC5AC-siRNA, AAV serotype 6 (AAV6) was selected for these studies based on its tropism for the lungs and ability to transduce airway epithelial cells *in vitro* and *in vivo*^27,28^. We have also shown in prior work that AAV6 can penetrate hyper-concentrated mucus produced by individuals with cystic fibrosis lung disease and transduce airway epithelium in the βENaC transgenic mouse model of muco-obstructive lung disease^29^. Building on this prior work, we determined in the current work that AAV6 can transduce secretory cells in mouse airways and as such, is a relevant gene vector for MUC5AC siRNA delivery. To determine its effectiveness in the context of disease, we further evaluated if AAV6 could successfully penetrate pathological asthma mucus for delivery of MUC5AC siRNA and improve MCC in IL-13 stimulated human airway epithelial cultures. The results of this work could open new avenues for inhaled therapeutics for asthma and other muco-obstructive lung diseases.

## Results

### AAV6 transduction in mouse airway and lung epithelium

AAV6 has shown tropism for airway and lung tissue with enhanced gene transfer efficiency *in vitro* and *in vivo* compared to other serotypes^28–31^. To examine and validate that AAV6 can deliver genes to target airway epithelial subtypes, we conducted an *in vivo* biodistribution study (**Fig 1**). Briefly, AAV6 encoding for an mCherry reporter was intranasally administered into 6-week-old BALB/c mice (5 males and 5 females) at a dose of 10^11^ viral particles/mouse in 15 µl PBS (**Fig 1A**) and equal volume of PBS in controls. As expected, mice in both the control and treated groups did not show any signs of adverse effects with normal weight gain observed over the course of the study (**Fig 1B**). After 14 days, mice were euthanized in order to collect trachea and lung tissue to measure mCherry expression in epithelial subtypes by flow cytometry and immunofluorescence. Immunofluorescence analysis showed mCherry expression in tracheal sections indicating successful AAV6 transduction (**Fig 1C**). Quantification of mCherry in tracheal epithelial subsets via flow cytometry indicated a significantly higher transduction in MUC5AC+ goblet cells subsets compared to basal and ciliated cells as well as increased transduction in TSPAN8+ secretory cells (**Fig 1D**). Immunofluorescence in lung samples yielded less conclusive results with low mCherry expression in immunofluorescence images of lung sections (**Fig S1A**). However, there was significantly higher transduction in TSPAN8+ epithelial cell subtypes compared to other epithelial subtypes (**Fig S1B**).

**Figure 1:**
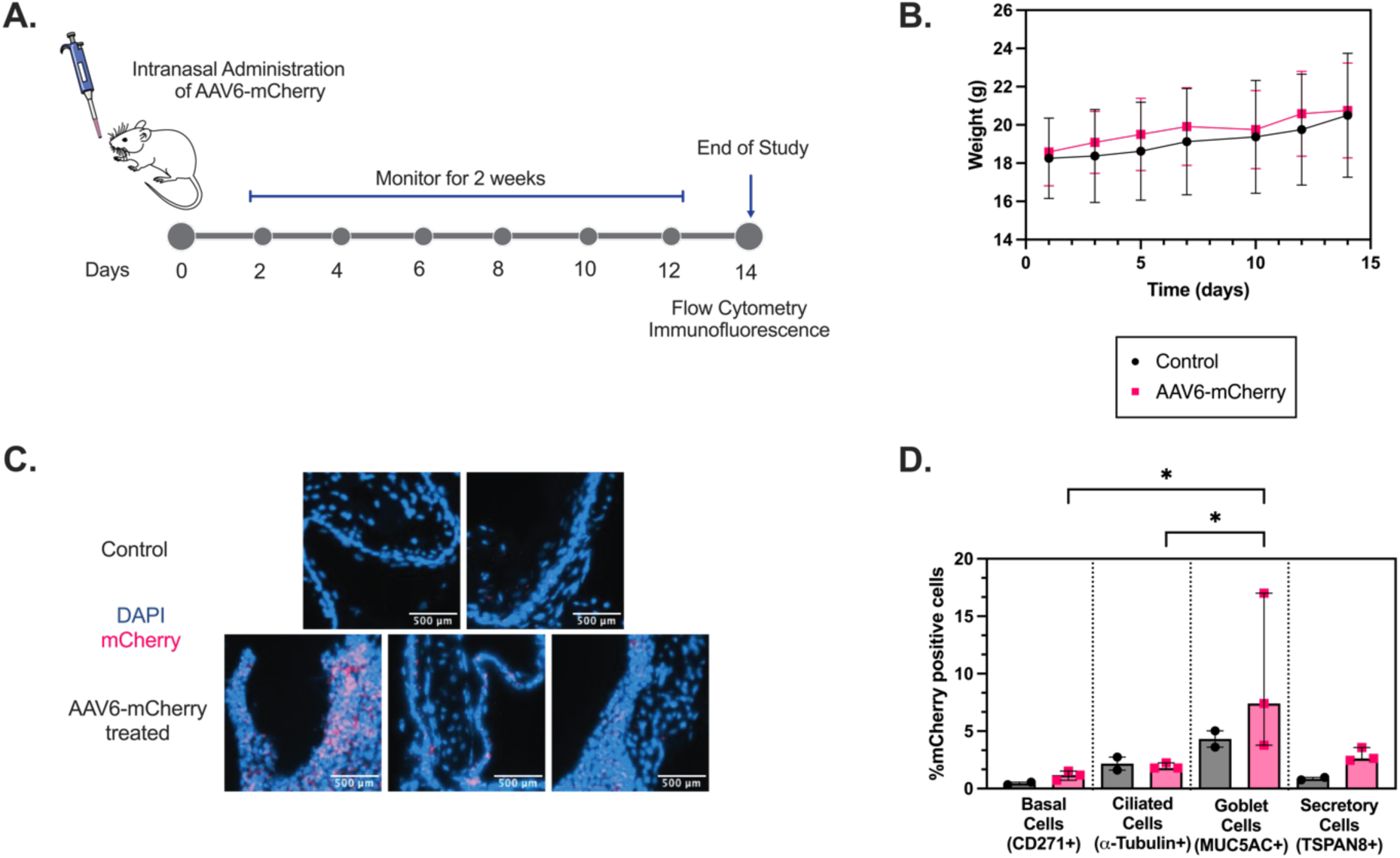
*In vivo* AAV6 transduction in mouse airway epithelial cells. **(A)** Schematic illustration showing intranasal AAV6 administration in BALB/c mice and overall study design. (B) Graph showing mouse weights which were recorded every other day throughout the study and showed a steady increase in both control and treated groups. **(C)** Immunofluorescence images of trachea sections in control (PBS) and AAV6-mCherry infected mice. mCherry transduction is shown in pink. Scale bar = 500 µm. **(D)** Bar Graphs show % mCherry positive cells normalized to control in different airway epithelial cells from single cell suspension of excised trachea. *\*p* < 0.05 by Ordinary one-way ANOVA. Each dot represents results from lung tissue from one mouse.

### AAV6 is diffusive in asthma-like, MUC5AC-enriched mucus

After confirming the relevance of AAV6 as a gene vector for pulmonary delivery to target mucus secreting cell types, we then characterized AAV6 diffusion in normal and MUC5AC-rich mucus produced *in vitro* from air liquid interface (ALI) cultures of human airway epithelial (HAE) cells to mimic mucus from asthmatic patients. Previous studies have shown that stimulation of HAE cultures with IL-13, which is a key cytokine in allergic asthma, causes goblet cell hyperplasia, increased MUC5AC production and reduced mucociliary transport^23,32–34^. As such, we established this *in vitro* asthma model by stimulating BCi-NS1.1 HAE cultures grown at ALI with IL-13 (10 ng/ml) for 7 days as previously described^22,33^ (**Fig 2A**). These cultures will be referred to as IL-13 treated HAE cultures. We also utilized a previously established model in BCi-NS1.1 cells where MUC5B gene expression is knocked-out via CRISPR-Cas9 leading to the production of MUC5AC-enriched airway mucus (**Fig 2A**).^35^ The diffusion of fluorescently labeled AAV6 (∼20 nm in diameter)^36^ and 20 nm diameter muco-inert nanoparticles (NP) was measured in control mucus. We confirmed that the muco-inert NPs were in the same size range as AAV6 by dynamic light scattering measurements. The diffusion rates for AAV6 and NPs were measured in the same regions of interest using fluorescence video microscopy. Based on the multiple particle tracking analysis and measured mean squared displacement at time scale of 1 second (MSD_1s_), the diffusion rate for muco-inert NPs were significantly higher than the rate for AAV6 (**Fig 2B**). We compared AAV6 diffusion in regular (healthy) mucus, IL-13 mucus, and mucus from MUC5B knockout (MUC5B KO) cultures, which are predominantly composed of MUC5AC to see how this change in mucus composition would impact viral diffusion. We found AAV6 is diffusive and traverses through the gel network comparably in control mucus, IL-13 mucus, and MUC5B-KO mucus as shown by the trajectories (**Fig 2C**) and median log_10_(MSD_1s_) values (**Fig 2D**) with no significant difference between mucus types.

**Figure 2:**
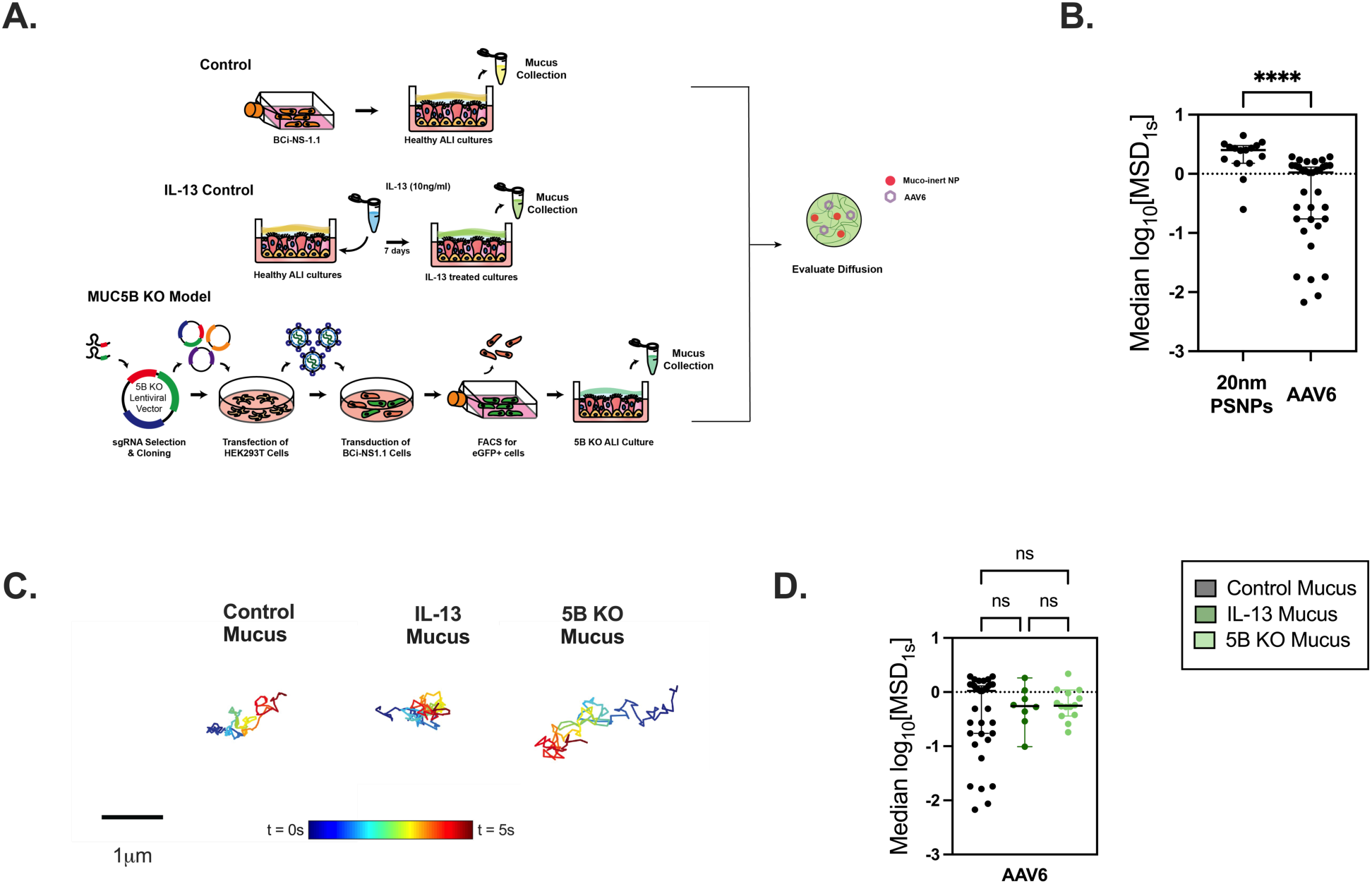
AAV6 diffusion in control and IL-13 treated mucus from *in vitro* cultures. **(A)** Schematic illustration showing the different *in vitro* models from which mucus was collected. BCi-NS1.1 cells were grown at ALI to establish Healthy ALI cultures. Healthy differentiated ALI cultures were stimulated with IL-13 (10 ng/ml) for 7 days to produce MUC5AC rich mucus. MUC5B KO cultures were generated using lentiviral mediated delivery of sgRNA and CRISPR-Cas9. Mucus from healthy controls, IL-13 treated cultures and MUC5B KO cultures were collected and AAV6 diffusion was measured in comparison to muco-inert nanoparticles of similar size. **(B)** Scatter plots of measured median log_10_[MSD_1s_] for AAV6 and 20 nm muco-inert nanoparticles in healthy mucus collected from ALI cultures. \*\*\*\**p* < 0.0001 by Mann-Whitney Test. **(C)** Representative trajectories of AAV6 diffusion in control mucus, IL-13 treated mucus, and MUC5B KO mucus. Trajectory color changes with time with dark blue indicating 0 s and dark red indicating 5 s. Scale bar = 1 µm. **(D)** Scatter plots of measured median log_10_[MSD_1s_] for AAV6 in healthy control mucus, IL-13 treated mucus, and MUC5B KO mucus. *p* > 0.05 by Kruskal-Wallis test with Dunn’s correction. Each dot represents data from 1 video (n≥3).

### AAV6 effectively transduces IL-13 stimulated airway epithelial cells in vitro

We next wanted to verify that AAV6 can transduce IL-13 stimulated HAE cultures *in vitro*. We first investigated the transduction efficiency of AAV6 expressing enhanced green fluorescence protein (eGFP) in undifferentiated BCi-NS1.1 HAE cells at multiplicities of infection (MOI) of 10^3^, 10^4^ and 10^5^. AAV6 mediated GFP expression was evaluated 72 hours post infection using fluorescence imaging. Successful transduction was observed in cultures inoculated at a MOI as low as 10^3^, with a substantial increase in fluorescence detection with MOI of 10^4^ and 10^5^ (**Fig 3A**). Based on this, AAV6-eGFP transduction was evaluated at MOI of 10^5^ in IL-13 treated differentiated ALI cultures and compared to untransduced untreated and IL-13 treated HAE controls. There was evident GFP expression in transduced groups (**Fig 3B**). The GFP integrated density was evaluated in AAV6-eGFP transduced ALI cultures with respect to controls (**Fig 3C**). There was a significantly higher expression of GFP in the transduced groups compared to controls.

**Figure 3:**
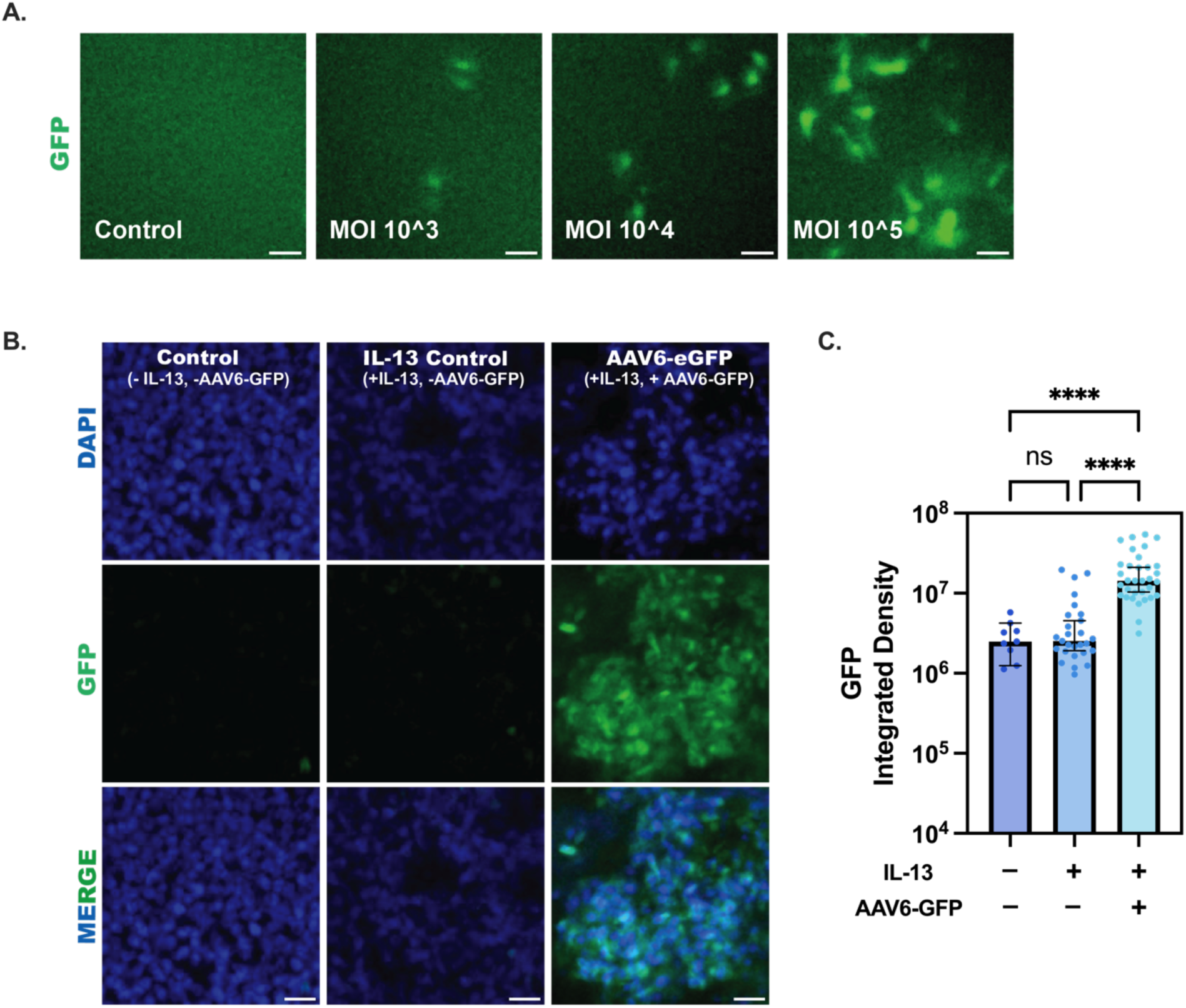
AAV6 transduction in healthy and IL-13 treated ALI cultures *in vitro*. **(A)** Representative images showing AAV6-eGFP transduction in undifferentiated BCi-NS1.1 cultures at MOI of 10^3, 10^4, and 10^5. Scale bar = 50 µm. **(B)** Representative images showing AAV6-eGFP transduction in IL-13 treated ALI cultures with respect to untransduced untreated and untransduced IL-13 treated controls. Scale bar = 20 µm. **(C)** Bar graphs showing GFP integrated density in IL-13 treated AAV6-eGFP transduced ALI cultures with respect to untransduced untreated and IL-13 treated controls (n=9). \*\*\**\*p* < 0.0001 by Kruskal-Wallis test with Dunn’s correction. Each dot represents different images.

### AAV6 mediated delivery of MUC5AC-siRNA in IL-13 stimulated human airway epithelial cells

AAVs can be effectively packaged with siRNA, which are short 21-nucleotide dsRNA molecules, for gene therapy to target disease-causing genes^37^. To test the efficacy of AAV6 mediated silencing of MUC5AC expression, we transduced ALI cultures with 2 doses of AAV6 carrying MUC5AC siRNA (AAV6-5AC siRNA) on day 15 and day 18 of differentiation respectively, at an MOI of 2500 and allowed 72h for transduction for each dose. On day 21, these cultures were challenged with IL-13 for 7 days. On day 28, the cultures were assessed for MUC5AC RNA and protein expression (**Fig 4A**). We performed qPCR to measure MUC5AC mRNA levels and found that groups treated with AAV6-5AC siRNA had comparable MUC5AC mRNA levels to untreated control whereas the IL-13 treated control was ∼1.5 times higher than the treated group and ∼2 times higher than untreated control (**Fig 4B**). To measure MUC5AC and MUC5B protein expression, we performed immunofluorescence staining in untreated, IL-13 treated control, and AAV6-5AC siRNA treated groups and quantified the integrated density of MUC5AC and MUC5B expression. There was increased MUC5AC expression in IL-13 treated controls compared to untreated control and AAV6-5AC siRNA treated groups (**Fig 4C & 4D**). There was also increased mucin content as reflected by the O-linked glycosylation quantification in IL-13 treated groups with lower and comparable mucin content in untreated control and AAV6-5AC siRNA treated group (**Fig S2**). MUC5B expression was comparable across all groups with no significant difference (**Fig 4C & 4E**).

**Figure 4:**
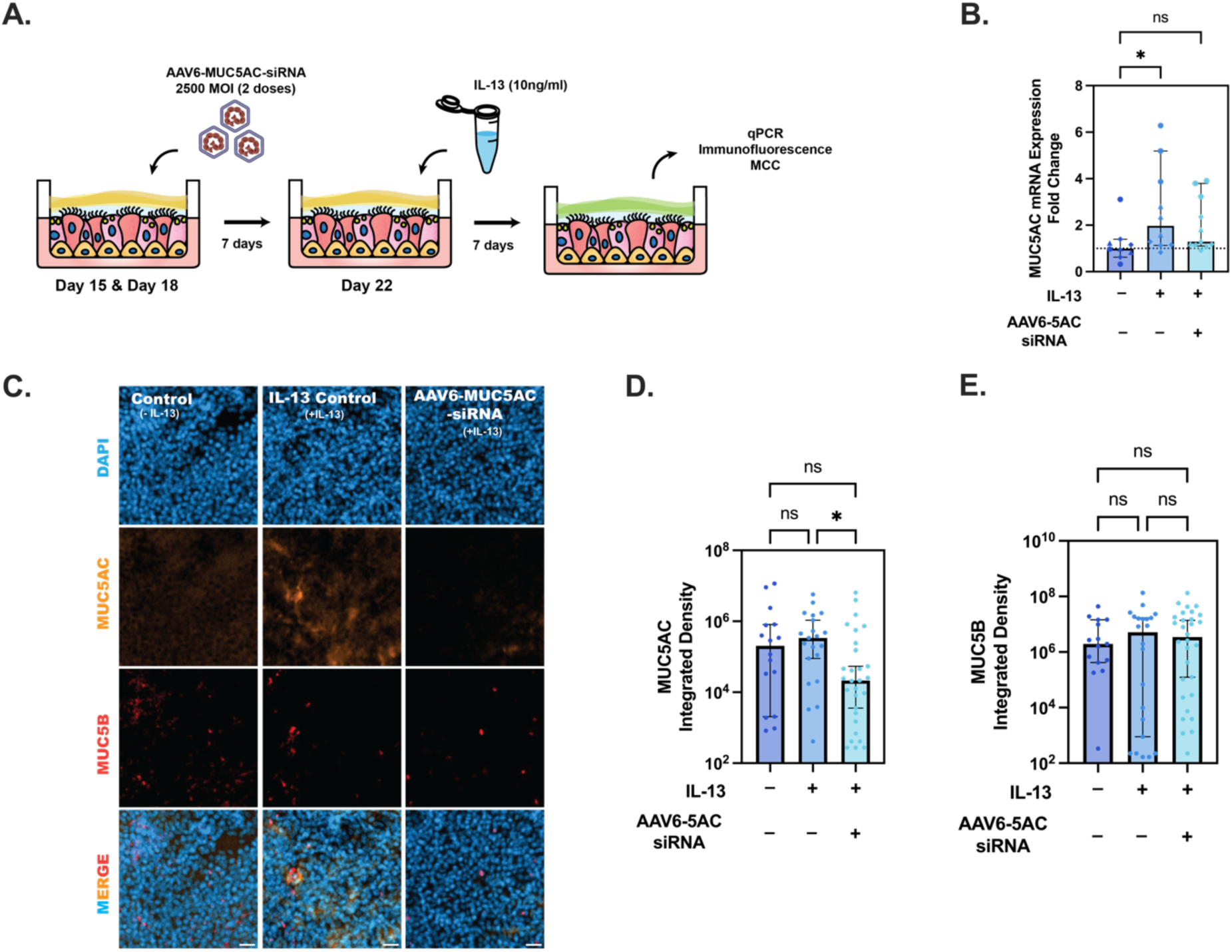
AAV6 delivers MUC5AC siRNA into IL-13 treated ALI cultures. **(A)** Schematic illustration demonstrating 2 dose pre-treatments with 2500 MOI of AAV6 expressing MUC5AC siRNA (AAV6-5AC siRNA) in ALI cultures before challenging with IL-13. **(B)** Scatter plot showing MUC5AC mRNA fold change in IL-13 treated control and AAV6-5AC siRNA treated groups with respect to untreated control. Shapes represent different experimental trials. *\*p* < 0.05 by Kruskal-Wallis test with Dunn’s correction. **(C)** Representative images showing MUC5B and MUC5AC protein expression in untreated control, IL-13 treated control and AAV6-5AC siRNA treated groups. Scale bar = 25 µm. **(D-E)** Bar graphs showing MUC5AC and MUC5B integrated density respectively, measured from immunofluorescence images (n≥15, across 9-15 cultures) in untreated controls, IL-13 treated controls, and AAV6-5AC siRNA treated groups. *\*p* < 0.05 by Kruskal-Wallis test with Dunn’s correction.

### Impact of AAV6 mediated MUC5AC siRNA treatment on mucociliary transport in IL-13 stimulated human airway epithelial cell cultures

To evaluate MUC5AC as a suitable target to improve airway clearance in asthma, we measured mucociliary transport rates following treatment with AAV6 expressing MUC5AC siRNA in IL-13 stimulated HAE cultures. Seven days after IL-13 challenge and/or AAV6-5AC siRNA treatment (**Fig 4A)**, mucociliary transport rate was measured by adding 2 µm red fluorescent beads to the apical surface of transwells and the movement of the beads were tracked in real time using live cell video microscopy (**Fig 5A**). We found that beads in IL-13 treated controls traversed less distance compared to beads on untreated controls and AAV6-5AC siRNA treated groups, which were comparable, as seen in representative trajectories (**Fig 5B**). We observed that the median transport rates of the beads in each video were comparable between untreated controls and AAV6-5AC siRNA treated groups and significantly reduced in IL-13 treated controls (**Fig 5C**). We next wanted to determine how many beads in each video were mobile. We defined beads to be mobile if they traversed a distance greater than or equal to their radii (1 µm) in 10 seconds equivalent to a velocity of 0.1 µm/s. Based on this, beads with speeds less than 0.1 µm/s were considered immobile or stuck (gray shaded region in **Fig 5D**). The percentage of mobile particle tracers in each video was then calculated using a 0.1 µm/s velocity cutoff. We found the untreated controls and AAV6-5AC siRNA treated groups had comparable mobile beads ∼65% while IL-13 treated controls had significantly fewer mobile beads, less than 50% were mobile (**Fig 5E**). To confirm that these changes in transport rates were due to change in secreted mucin composition and not ciliated epithelium function, we measured the ciliary beat frequency which was comparable in all groups (**Fig 5F**).

**Figure 5:**
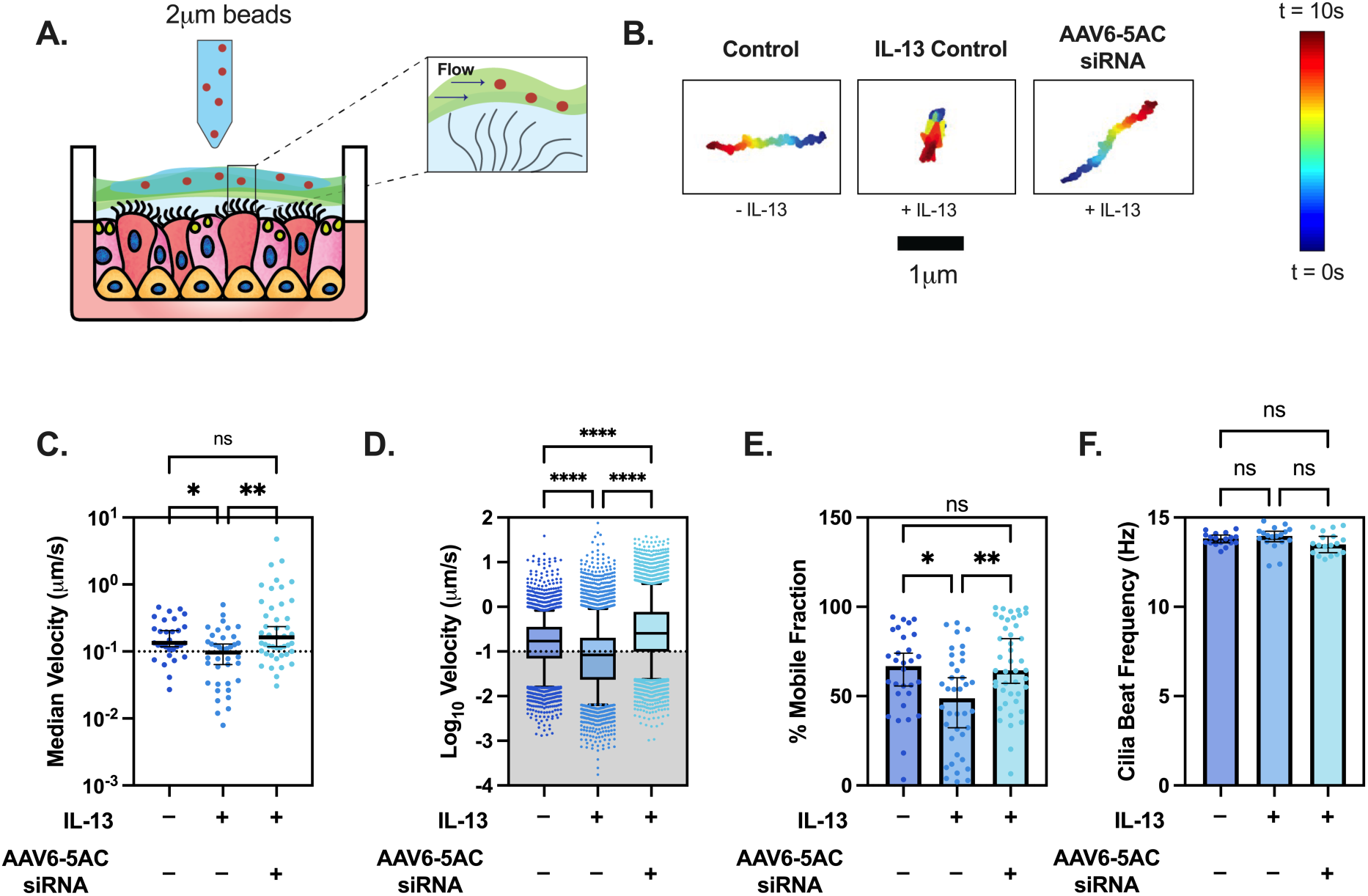
Mucociliary transport is maintained in MUC5AC siRNA treated HAE cultures challenged with IL-13. **(A)** Schematic showing the design of MCT measurement. Briefly, 2 µm beads are applied onto mucus on ALI cultures. The coordinated beating of cilia causes the mucus with beads to flow, which is imaged. **(B)** Representative trajectories of single beads in untreated control, IL-13 treated control, and AAV6-5AC siRNA treated group. Dark blue represents trajectories at 0 s and dark red represents trajectories at 10 s. (**C**) Scatter plot represents median velocities (µm/s) of beads in each video for untreated control, IL-13 treated control, and AAV6-5AC siRNA groups (n = 30). *\*\*p* < 0.01 by Kruskal-Wallis test with Dunn’s correction. (**D**) Box and Whisker plot representing velocities (µm/s) of all beads in untreated control, IL-13 treated control and AAV6-5AC siRNA groups. Shaded region indicates particles with velocities < 0.1 µm/s. *\*\*\*\*p* < 0.0001 by Kruskal-Wallis test with Dunn’s correction. (**E**) Scatter plot showing immobile fraction (no. of beads with velocity < 0.1 µm/s/ total beads) in each video for untreated control, IL-13 treated control and AAV6-5AC siRNA groups. *\*\*\*p* < 0.001 by Kruskal-Wallis test with Dunn’s correction. (**F**) Bar graph showing ciliary beat frequency (Hz) in untreated control, IL-13 treated control, and AAV6-5AC siRNA groups.

## Discussion

In these studies, we targeted the mucin glycoprotein MUC5AC using AAV vectors to explore its potential as an inhaled gene therapy for asthma. We first assessed the capabilities of AAV6 to successfully deliver cargo into mouse lungs and airways via intranasal administration and determine if relevant epithelial subsets were transduced (e.g. mucus-secreting cells). Previous studies have shown successful AAV6 transduction of airway epithelial cells^27–29^, but transduction in epithelial subsets have not been analyzed. We saw that AAV6 successfully transduced mucus secreting airway epithelial cells in the trachea. Overall, transduction in the lungs was minimal, which is to be expected for the intranasal route of administration. We anticipate higher transduction in the deeper lung via intratracheal modes of administration^31^ which will be tested in subsequent *in vivo* efficacy studies. Detection of AAV6 transduction in goblet cells in the trachea is promising and provides further rationale for using AAV6 to deliver siRNA targeting MUC5AC in asthma.

A significant biological barrier to inhaled gene delivery vectors is the pathological mucus produced in diseases like asthma. We found in prior work that AAV6 can diffuse through pathological mucus from individuals with CF and transduce airway epithelial cells *in vivo*^29^. Similarly, we saw that AAV6 was equally diffusive in pathological *in vitro* asthma-like (i.e.., IL-13 stimulated) mucus, normal mucus, and MUC5B KO (i.e., MUC5AC-rich) mucus collected from ALI cultures. AAV6 diffusion was lower than muco-inert nanoparticles which may possibly be due to interactions of AAV6 capsid proteins with heparan sulfate terminated glycans present in airway mucus^38,39^. However, overall, AAV6 retained the capacity to transduce the underlying epithelium in IL-13 stimulated, MUC5AC-secreting HAE cultures.

Pathological viscous asthma mucus is the consequence of increased goblet cell number and therefore increased MUC5AC expression^21,40^. Consistent with prior studies, we observed increased mRNA and protein expression of MUC5AC in IL-13 treated cultures. Pre-treatment with AAV6 expressing MUC5AC siRNA was able to protect cultures when challenged with IL-13, with MUC5AC expression comparable to untreated controls. This was also reflected in the quantification of total O-linked glycosylation in mucus collected from each group. In terms of MUC5B expression, prior literature is inconsistent, with some studies reporting increased MUC5B expression in asthma^41^, and others reporting no change^42^. In the current work, we observed no significant differences in MUC5B protein expression between the groups and as such, MUC5AC siRNA treatment did not appear to cause any compensatory expression of MUC5B.

Impaired mucociliary clearance in asthma contributes to obstruction in the airway and creates a niche for exacerbation-initiating infections^19,43^. As noted, asphyxiation from complete blocking of the airway by intraluminal mucus plugs is the primary cause of death in asthma^44–46^. To determine the potential of the approach to restore normal MCC in asthma, we also evaluated if MUC5AC siRNA delivery with AAV6 would lead to improved MCC in IL-13 stimulated HAE cultures. In IL-13 treated HAE cultures pre-treated with AAV6-5AC siRNA, MCC remained effective despite being challenged with IL-13 with comparable transport velocities and mobile beads to untreated control. These findings suggests a prophylactic strategy to suppress MUC5AC hypersecretion could be useful in preventing mucostasis and generation of airway mucus plugs in asthma.

The present study demonstrated that AAV6 is successfully able to penetrate an asthma-like mucus barrier and transduce IL-13 stimulated HAE cells differentiated at ALI *in vitro*. We also showed using an *in vitro* model of IL-13 induced asthma that AAV6 mediated siRNA delivery could effectively prevent MUC5AC hypersecretion and mucociliary dysfunction. Our study also provides rationale for development of mucin-targeted gene therapies for asthma and potentially other chronic respiratory diseases using AAV6 given its capability to transduce secretory cells *in vivo*. These studies provide the basis for our future plans to evaluate the effectiveness of AAV6 mediated MUC5AC siRNA treatment in an *in vivo* model of allergic asthma. A limitation of the current work is this approach was only tested as a prophylactic treatment prior to establishment of inflammation and mucostasis with IL-13 stimulation. Therefore, we also plan to assess if MUC5AC siRNA delivery has the potential to reverse MUC5AC hypersecretion *in vitro* and *in vivo.* Our results establish AAV gene therapy for MUC5AC-targeting siRNA delivery as an attractive option to prevent mucus obstruction as a treatment for asthma.

## Materials and Methods

### Cell Culture

The BCi-NS1.1 immortalized human basal cell line was generously provided by Ronald Crystal (Weill Cornell Medical College) and cultured based on previous methods^35,47^. MUC5B KO BCi-NS1.1 cultures were generated as previously reported^35^ and cultures were prepared from frozen stocks for experiments. BCi-NS1.1 cells were cultured at 37°C, 5% CO_2_ in a flask with PneumaCult-Ex Plus expansion media (STEMCELL Technologies). At about 80% confluency the cells were detached using 0.05% Trypsin-EDTA for 5 mins and seeded on 12 mm rat-tail collagen type-I coated transwell inserts (StemCell Technologies) at 10,000 cells/cm^2^ in Ex Plus medium supplied on both the apical and basolateral compartments, until a confluent monolayer was established. Once confluent, medium from the apical compartment was removed and PneumaCult-ALI media (STEMCELL Technologies) was supplied only to the basolateral compartment to transition the cultures into Air-Liquid Interface cultures. The cells were allowed to differentiate into a pseudostratified mucociliary epithelium for 4 weeks with media changed every other day. To establish an *in vitro* asthma model, ALI cultures were treated with recombinant human IL-13 (10 ng/ml, Peprotech) added in the basolateral compartment for 7 days.

### Mucus collection

Mucus was collected from fully differentiated ALI cultures once or twice a week. PBS was added to the apical surface of cultures for 30 mins at 37°C. After 30 mins, mucus washings were concentrated by passing through 100kDa amicon filter and were stored at -80°C for long term storage prior to usage.

### Particle Tracking Microrheology (PTM)

Based on previously reported methods^48^, carboxylate-modified polystyrene nanoparticles (ThermoFisher) were surface modified with a 5-kDa polyethylene glycol-amine (PEG) coating to ensure that these particles are non-adhesive to mucus. Size and zeta potential of the particles were measured using NanoBrook Omni (Brookhaven Instruments). The nanoparticles had a hydrodynamic diameter of 27.74 ± 10.99 nm. Twenty microliters of mucus was added to the microscopy chamber made from vacuum-grease coated O-rings in which 1 µl of nanoparticles and fluorescently labeled AAV6 respectively (∼0.002% w/v) were added and allowed to incubate for 30 mins at room temperature. The diffusion of nanoparticles and AAV6 was measured using fluorescence video imaging using Zeiss 800 LSM Microscope at 63x-water immersion objective. Briefly, videos were taken for 10 s at 300ms exposure time (30 Hz) and analyzed using previously developed MATLAB algorithm^49^. The mean squared displacement (MSD) for each particle for lag time (r) was calculated as (MSD(r)) = ([*x*(*t*+r)–*x*(*t*)]^2^ + [*y*(*t*+r)–y(*t*)]^2^).

### AAV6-eGFP Transduction In Vitro

Undifferentiated BCi-NS1.1 cultures were seeded in 6 well plates. At 60-65% confluency, the cells were infected with AAV6 encoding GFP (AAV6-CAG-eGFP, 10^13^ viral particles/ml, AAVner Gene) at multiplicities of infection (MOI) of 1000, 10,000, and 100,000 viral particles per cell. The cells were imaged on a Zeiss 800 LSM Microscope at 72 hours post infection and analyzed for GFP expression. Differentiated BCi-NS1.1 cultures were transduced with AAV6-eGFP at MOI 100,000 apically in regular and IL-13 stimulated cultures, and the cells were fixed and imaged for GFP expression after 72 hours.

### AAV6 MUC5AC siRNA Transduction In Vitro

AAV6 packaged with MUC5AC siRNA (NM_001304359) was obtained from Applied Biological Materials, Canada. Briefly, ALI cultures were pre-treated apically with 2500 MOI on with their 1^st^ dose on day 15 of differentiation and a second dose on day 18 of differentiation. These cultures were challenged with IL-13 (10 ng/ml) supplemented to the media in the basolateral chamber from day 21-28 and replenished with every media change. On day 28, the cells were analyzed using PCR, immunofluorescent staining and particle velocimetry measurements of MCC.

### Immunofluorescence

Differentiated and undifferentiated BCi-NS1.1 cells were fixed with 100% ice-cold methanol for 15 mins at 4°C. The cells were washed with PBS and blocked with 5% bovine serum albumin (BSA) in 0.01% PBST for 1 hour at room temperature. For MUC5B (rabbit anti-human, Cell Signaling Technologies, MA) and MUC5AC (mouse anti-human, ThermoFisher Scientific) expression, the cultures were incubated with primary antibodies (1:1000) overnight at 4°C. The next day, cultures were washed with PBS twice. The cultures were incubated with secondary antibodies AlexaFlour 647 (anti-rabbit, 1:2000) and AlexaFlour 555 (anti-mouse, 1:2000) in PBS for 1 hour at room temperature. The cultures were then washed with PBS twice. For cultures with AAV6-eGFP transduction, cultures were fixed, incubated in 1 µg/ml DAPI in PBS for 15 mins at room temperature, washed twice with PBS, and then mounted and imaged using Zeiss 800 LSM Microscope. Quantification of integrated density of MUC5B and MUC5AC expression was measured in FIJI image software using the automated thresholding function.

### qRT-PCR

RNA was extracted from differentiated BCi-NS1.1 cultures using the RNeasy minikit (Qiagen) based on manufacturer’s protocol. cDNA from the RNA was prepared with SuperScript III (Invitrogen) as per manufacturer’s protocol. Quantitative PCR for MUC5AC was carried out using PowerUP SYBR Green Master Mix (Applied Biosystems) using forward and reverse primers (Integrated DNA Technologies).

### Mucociliary transport and ciliary beat frequency

Prior to MCC and CBF analyses, cultures were washed, and mucus was allowed to accumulate for 24 hours. A 25 µl suspension of 2 µm red-fluorescent polystyrene microspheres (Invitrogen, 1:2000 dilution in PBS) was applied apically to the cultures and allowed to equilibrate at 37°C for 15 mins. Videos of 5 regions were recorded using the Zeiss 800 LSM Microscope at 10x at frame rate of 10 Hz for 10 seconds. The microsphere tracking analysis is based on a custom MATLAB algorithm^22^. The software computes the trajectories of the fluorescent microsphere in the xy-plane in each frame. The displacement is calculated based on trajectories and velocity is calculated based on displacement over time elapsed. To measure ciliary beat frequency, 20 second videos at frame rate of 50 Hz were recorded at 10x magnification using the Zeiss 800 LSM Microscope from 3 random regions using brightfield. The local pixel intensity maxima were counted using custom written MATLAB algorithm, which indicates beating cilia. The beat frequency was calculated by dividing the number of beats over time elapsed.

### AAV6-mCherry Transduction In Vivo

Six-week-old male and female BALB/c mice were purchased from Charles River Laboratories (Wilmington, MA) and maintained in disease free conditions at the animal facility. The animal study protocol was approved by the Institutional Animal Care and Use Committee (IACUC) of the University of Maryland College Park (protocol R-MAY-22-25). AAV6 encoding mCherry (AAV6-CMV-mCherry, stock: 10^13 viral particles/ml, AAVner Gene) was intranasally administered to anesthetized mice at a dose of 10^11^ viral particles/mouse. The mice were monitored for 2 weeks and were sacrificed on day 14. Their lungs and trachea were isolated for imaging and flow cytometry analysis as described below.

### Flow Cytometry

The lungs and tracheas of mice were removed and dissected on petri plates. Single cell suspension of the tissues was made by digesting tissue with Collagenase IV (1mg/ml, ThermoFisher Scientific), and DNAse I (25ug/ml, Millipore Sigma) for 45 minutes at 37°C. Dispase II (2mg/ml, Millipore Sigma) was also added for trachea. The tissue was then repeatedly passed through a 70 µm cell strainer until a single cell suspension was obtained. The cells were washed in PBS (300xg, 5 mins) twice. The lung cells were suspended in RBC Lysis buffer for 3 mins on ice and washed with PBS. The cells were distributed into groups in FACS staining buffer and were stained for different airway epithelial cell markers as shown in **Table 1** and flow cytometry was performed (BD FACS Celesta).

**Table 1.**
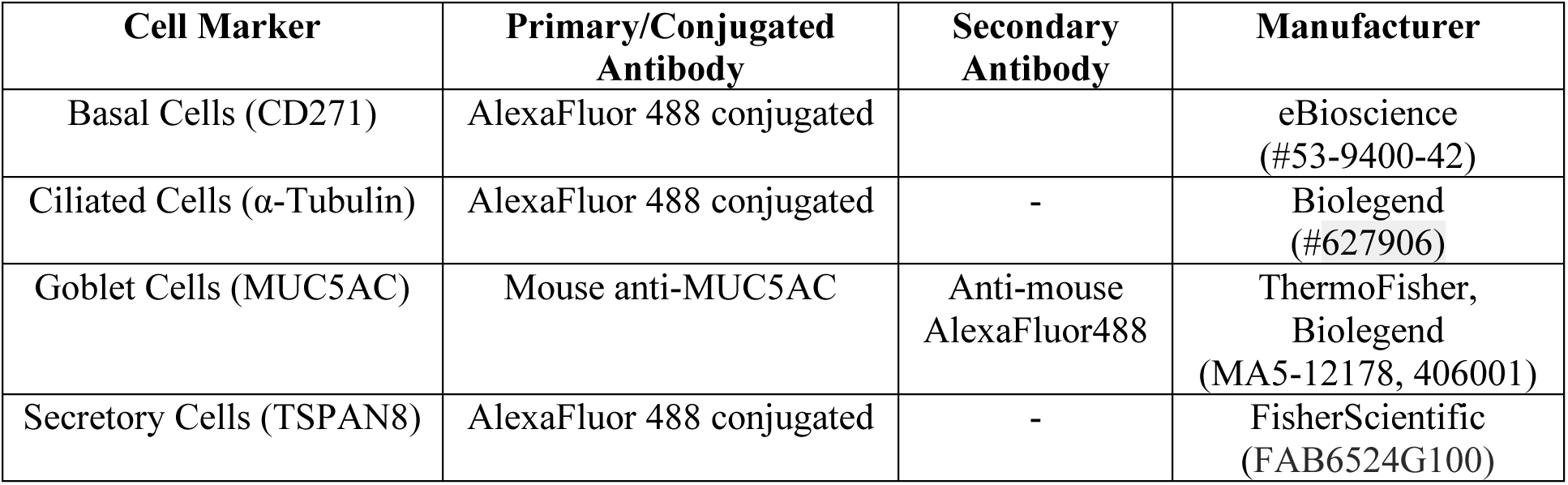
Antibodies used for flow cytometry.

| Cell Marker | Primary/Conjugated Antibody | Secondary Antibody | Manufacturer |
| --- | --- | --- | --- |
| Basal Cells (CD271) | AlexaFluor 488 conjugated |  | eBioscience (#53-9400-42) |
| Ciliated Cells (α-Tubulin) | AlexaFluor 488 conjugated | - | Biolegend (#627906) |
| Goblet Cells (MUC5AC) | Mouse anti-MUC5AC | Anti-mouse AlexaFluor488 | ThermoFisher, Biolegend (MA5-12178, 406001) |
| Secretory Cells (TSPAN8) | AlexaFluor 488 conjugated | - | FisherScientific (FAB6524G100) |

### Ex Vivo Cryosectioning and Staining

To prepare lungs and trachea for immunofluorescence, the lungs were inflated with 10% formalin which was delivered through blunt-end needle inserted via the trachea. After inflation, the lungs and trachea were collected and placed in 10% formalin overnight at room temperature. The tissues were transferred to cryomolds with OCT and frozen at -80°C. Using the Leica CM1950 Cryostat, 10 µm lung sections and 7 µm tracheal sections were made. The sections were allowed to calibrate at room temperature overnight and fixed with 100% methanol and stained with DAPI as per immunofluorescence protocol and imaged at 10x magnification using the Zeiss 800 LSM Microscope.

### Statistical Analysis

The data were statistically analyzed using GraphPad Prism Software. Comparison between 2 groups were performed using 2-tailed student *t* test or Mann-Whitney *U* test. For comparison between groups, either one-way analysis of variance (ANOVA) followed by a Tukey post hoc correction was used or Kruskal-Wallis with Dunn’s correction was used for non-Gaussian distributions. All graphs show median values and 5th up to 95th percentiles of data and outliers are included. Differences were considered statistically significant at *p* value < 0.05.

## Funding

This project was funded by National Institutes of Health (R01 HL160540) and University of Maryland Grand Challenges Research Grant (GC18).

## Author contributions

Conceptualization: GAD

Methodology: SK, YC, MC, MAS, GAD

Investigation: SK, YC, MC

Visualization: SK, GAD

Supervision: MAS, GAD

Writing—original draft: SK

Writing—review & editing: SK, YC, MC, MAS, GAD

## Competing interests

Authors declare that they have no competing interests.

## Data and materials availability

All data are available in the main text or the supplementary materials.

## Notes

### Competing Interest Statement

The authors have declared no competing interest.

## References

1. Braman, S. S. The Global Burden of Asthma. Chest 130, 4S–12S (2006).

2. Dharmage, S. C., Perret, J. L. & Custovic, A. Epidemiology of Asthma in Children and Adults. Frontiers in Pediatrics 7, (2019).

3. The Global Asthma Report 2022. int j tuberc lung dis 26, 1–104 (2022).

4. Fahy, J. V. Type 2 inflammation in asthma--present in most, absent in many. Nat Rev Immunol 15, 57–65 (2015).

5. Spellberg, B. & Edwards, J. E. Type 1/Type 2 immunity in infectious diseases. Clin Infect Dis 32, 76–102 (2001).

6. Song, D., Cahn, D. & Duncan, G. A. Mucin Biopolymers and Their Barrier Function at Airway Surfaces. Langmuir 36, 12773–12783 (2020).

7. Patel, V. H. et al. Current Limitations and Recent Advances in the Management of Asthma. Disease-a-Month 69, 101483 (2023).

8. Ip, M., Lam, K., Yam, L., Kung, A. & Ng, M. Decreased bone mineral density in premenopausal asthma patients receiving long-term inhaled steroids. Chest 105, 1722–1727 (1994).

9. Gaga, M. & Zervas, E. Oral steroids in asthma: a double-edged sword. Eur Respir J 54, 1902034 (2019).

10. Boboltz, A., Kumar, S. & Duncan, G. A. Inhaled drug delivery for the targeted treatment of asthma. Advanced Drug Delivery Reviews 198, 114858 (2023).

11. Zavorotinskaya, T., Tomkinson, A. & Murphy, J. E. Treatment of experimental asthma by long-term gene therapy directed against IL-4 and IL-13. Molecular Therapy 7, 155–162 (2003).

12. Behera, A. K., Kumar, M., Lockey, R. F. & Mohapatra, S. S. Adenovirus-Mediated Interferon γ Gene Therapy for Allergic Asthma: Involvement of Interleukin 12 and STAT4 Signaling. Human Gene Therapy 13, 1697–1709 (2002).

13. Xie, Y. et al. Targeted delivery of siRNA to activated T cells via transferrin-polyethylenimine (Tf-PEI) as a potential therapy of asthma. Journal of Controlled Release 229, 120–129 (2016).

14. Choi, M., Gu, J., Lee, M. & Rhim, T. A new combination therapy for asthma using dual-function dexamethasone-conjugated polyethylenimine and vitamin D binding protein siRNA. Gene Ther 24, 727–734 (2017).

15. Thanki, K., Blum, K. G., Thakur, A., Rose, F. & Foged, C. Formulation of RNA interference-based drugs for pulmonary delivery: challenges and opportunities. Therapeutic Delivery 9, 731–749 (2018).

16. Tanaka, M. & Nyce, J. W. Respirable antisense oligonucleotides: a new drug class for respiratory disease. Respir Res 2, 5–9 (2001).

17. Chiang, P.-C., Chen, J.-C., Chen, L.-C. & Kuo, M.-L. Adeno-Associated Virus-Mediated Interleukin-12 Gene Expression Alleviates Lung Inflammation and Type 2 T-Helper-Responses in Ovalbumin-Sensitized Asthmatic Mice. Human Gene Therapy 33, 1052–1061 (2022).

18. Dunican, E. M., Watchorn, D. C. & Fahy, J. V. Autopsy and Imaging Studies of Mucus in Asthma. Lessons Learned about Disease Mechanisms and the Role of Mucus in Airflow Obstruction. Ann Am Thorac Soc 15, S184–S191 (2018).

19. Dunnill, M. S. THE PATHOLOGY OF ASTHMA, WITH SPECIAL REFERENCE TO CHANGES IN THE BRONCHIAL MUCOSA. J Clin Pathol 13, 27–33 (1960).

20. Huber, H. L. & Koessler, K. K. THE PATHOLOGY OF BRONCHIAL ASTHMA. Archives of Internal Medicine 30, 689–760 (1922).

21. Lachowicz-Scroggins, M. E. et al. Abnormalities in MUC5AC and MUC5B Protein in Airway Mucus in Asthma. Am J Respir Crit Care Med 194, 1296–1299 (2016).

22. Song, D. et al. Modeling Airway Dysfunction in Asthma Using Synthetic Mucus Biomaterials. ACS Biomaterials Science & Engineering (2021) doi:10.1021/acsbiomaterials.0c01728.

23. Bonser, L. R., Zlock, L., Finkbeiner, W. & Erle, D. J. Epithelial tethering of MUC5AC-rich mucus impairs mucociliary transport in asthma. Journal of Clinical Investigation 126, 2367–2371 (2016).

24. Siddiqui, S., et al. Epithelial miR-141 regulates IL-13–induced airway mucus production. JCI Insight 6, e139019 (2021).

25. Wang, J.-H., Gessler, D. J., Zhan, W., Gallagher, T. L. & Gao, G. Adeno-associated virus as a delivery vector for gene therapy of human diseases. Sig Transduct Target Ther 9, 1–33 (2024).

26. Liu, D. et al. Adeno-associated virus therapies: Pioneering solutions for human genetic diseases. Cytokine & Growth Factor Reviews 80, 109–120 (2024).

27. Halbert, C. L., Allen, J. M. & Miller, A. D. Adeno-Associated Virus Type 6 (AAV6) Vectors Mediate Efficient Transduction of Airway Epithelial Cells in Mouse Lungs Compared to That of AAV2 Vectors. Journal of Virology 75, 6615–6624 (2001).

28. Li, W., Zhang, L., Wu, Z., Pickles, R. J. & Samulski, R. J. AAV-6 mediated efficient transduction of mouse lower airways. Virology 417, 327–333 (2011).

29. Duncan, G. A. et al. An adeno-associated viral vector capable of penetrating the mucus barrier to inhaled gene therapy. Molecular Therapy - Methods & Clinical Development 9, 296–304 (2018).

30. Limberis, M. P., Vandenberghe, L. H., Zhang, L., Pickles, R. J. & Wilson, J. M. Transduction Efficiencies of Novel AAV Vectors in Mouse Airway Epithelium In Vivo and Human Ciliated Airway Epithelium In Vitro. Mol Ther 17, 294–301 (2009).

31. Kurosaki, F. et al. Optimization of adeno-associated virus vector-mediated gene transfer to the respiratory tract. Gene Ther 24, 290–297 (2017).

32. Turner, J. et al. Goblet cells are derived from a FOXJ1-expressing progenitor in a human airway epithelium. Am J Respir Cell Mol Biol 44, 276–284 (2011).

33. Laoukili, J. et al. IL-13 alters mucociliary differentiation and ciliary beating of human respiratory epithelial cells. J Clin Invest 108, 1817–1824 (2001).

34. Koh, K. D. et al. Efficient RNP-directed Human Gene Targeting Reveals SPDEF Is Required for IL-13-induced Mucostasis. Am J Respir Cell Mol Biol 62, 373–381 (2020).

35. Song, D. et al. MUC5B mobilizes and MUC5AC spatially aligns mucociliary transport on human airway epithelium. Science Advances 8, eabq5049 (2022).

36. Liu, X. et al. Evaluation of a rapid multi-attribute combinatorial high-throughput UV-Vis/DLS/SLS analytical platform for rAAV quantification and characterization. Molecular Therapy Methods & Clinical Development 32, (2024).

37. Tomar, R. S., Matta, H. & Chaudhary, P. M. Use of adeno-associated viral vector for delivery of small interfering RNA. Oncogene 22, 5712–5715 (2003).

38. Wu, Z. et al. Single Amino Acid Changes Can Influence Titer, Heparin Binding, and Tissue Tropism in Different Adeno-Associated Virus Serotypes. J Virol 80, 11393–11397 (2006).

39. Xie, Q., Lerch, T. F., Meyer, N. L. & Chapman, M. S. Structure-function analysis of receptor-binding in adeno-associated virus serotype 6 (AAV-6). Virology 420, 10–19 (2011).

40. Bonser, L. R. & Erle, D. J. Airway Mucus and Asthma: The Role of MUC5AC and MUC5B. J Clin Med 6, 112 (2017).

41. Kirkham, S., Sheehan, J. K., Knight, D., Richardson, P. S. & Thornton, D. J. Heterogeneity of airways mucus: variations in the amounts and glycoforms of the major oligomeric mucins MUC5AC and MUC5B. Biochem J 361, 537–546 (2002).

42. Welsh, K. G. et al. MUC5AC and a Glycosylated Variant of MUC5B Alter Mucin Composition in Children With Acute Asthma. Chest 152, 771–779 (2017).

43. Button, B. et al. Periciliary Brush Promotes the Lung Health by Separating the Mucus Layer from Airway Epithelia. Science 337, 937–941 (2012).

44. Messer, J. W., Peters, G. A. & Bennett, W. A. Causes of death and pathologic findings in 304 cases of bronchial asthma. Dis Chest 38, 616–624 (1960).

45. Kuyper, L. M. et al. Characterization of airway plugging in fatal asthma. Am J Med 115, 6–11 (2003).

46. Evans, C. M., Kim, K., Tuvim, M. J. & Dickey, B. F. Mucus hypersecretion in asthma: causes and effects. Curr Opin Pulm Med 15, 4–11 (2009).

47. Walters, M. S. et al. Generation of a human airway epithelium derived basal cell line with multipotent differentiation capacity. Respir Res 14, 135 (2013).

48. Schuster, B. S., Suk, J. S., Woodworth, G. F. & Hanes, J. Nanoparticle diffusion in respiratory mucus from humans without lung disease. Biomaterials 34, 3439–3446 (2013).

49. Schuster, B. S., Ensign, L. M., Allan, D. B., Suk, J. S. & Hanes, J. Particle tracking in drug and gene delivery research: state-of-the-art applications and methods. Adv Drug Deliv Rev 91, 70–91 (2015).

