## Supporting Information for "AAV-mediated MUC5AC siRNA delivery to prevent mucociliary dysfunction in asthma"

**Affiliations**

**Supplementary Information**

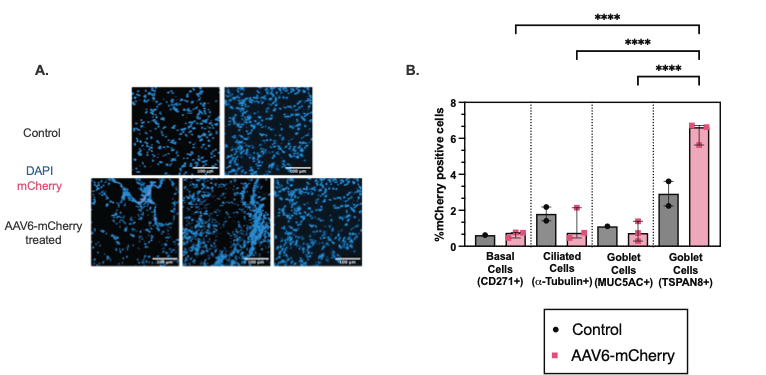

**Figure S1: *In vivo* AAV6 transduction in mouse lungs. (A)** Immunofluorescence images of lung sections in control (PBS) and AAV6-mCherry infected mice. mCherry transduction is shown in pink. Scale bar = 100 µm. **(B)** Bar Graphs show % mCherry positive cells normalized to control in different airway epithelial cells from single cell suspension of excised trachea. *****p* < 0.0001 by Ordinary one-way ANOVA. Each dot represents a mouse.

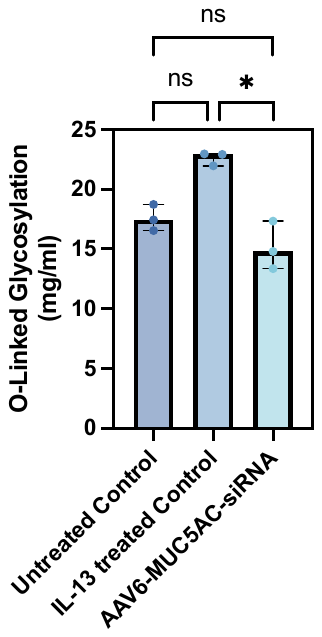

**Figure S2: *In vitro* mucin content.** Bar graphs showing O-Linked glycosylation in controls and treated groups. **p* < 0.05 by Kruskal-Wallis test with Dunn’s correction.
